# Heterogeneous expression of the SARS-Coronavirus-2 receptor ACE2 in the human respiratory tract

**DOI:** 10.1101/2020.04.22.056127

**Authors:** Miguel E. Ortiz Bezara, Andrew Thurman, Alejandro A. Pezzulo, Mariah R. Leidinger, Julia A. Klesney-Tait, Philip H. Karp, Ping Tan, Christine Wohlford-Lenane, Paul B. McCray, David K. Meyerholz

## Abstract

**Background:** Zoonotically transmitted coronaviruses are responsible for three disease outbreaks since 2002, including the current COVID-19 pandemic, caused by SARS-CoV-2. Its efficient transmission and range of disease severity raise questions regarding the contributions of virus-receptor interactions. ACE2 is a host ectopeptidase and the receptor for SARS-CoV-2. Numerous reports describe ACE2 mRNA abundance and tissue distribution; however, mRNA abundance is not always representative of protein levels. Currently, there is limited data evaluating ACE2 protein and its correlation with other SARS-CoV-2 susceptibility factors.

**Materials and methods:** We systematically examined the human upper and lower respiratory tract using single-cell RNA sequencing and immunohistochemistry to determine receptor expression and evaluated its association with risk factors for severe COVID-19.

**Findings:** Our results reveal that ACE2 protein is highest within regions of the sinonasal cavity and pulmonary alveoli, sites of presumptive viral transmission and severe disease development, respectively. In the lung parenchyma, ACE2 protein was found on the apical surface of a small subset of alveolar type II cells and colocalized with TMPRSS2, a cofactor for SARS-CoV2 entry. ACE2 protein was not increased by pulmonary risk factors for severe COVID-19.

Additionally, ACE2 protein was not reduced in children, a demographic with a lower incidence of severe COVID-19.

**Interpretation:** These results offer new insights into ACE2 protein localization in the human respiratory tract and its relationship with susceptibility factors to COVID-19.

**Research in context:** *Evidence before this study:* Previous studies of ACE2 mRNA transcript abundance in the human respiratory tract have suggested a possible association between ACE2 expression and age, sex, and the presence of comorbidities. However, these studies have provided conflicting results, as well as a lack of protein validation. Previous ACE2 protein studies have been limited by a paucity of lung tissue samples and reports that have produced contradictory results.

*Added value of this study:* Using a combination of single-cell RNA sequencing and immunohistochemistry, we describe ACE2 expression in the human respiratory tract. Staining protocols were optimized and validated to show consistent apical localization and avoid non-specific staining. We show ACE2 protein is found in subsets of airway cells and is highest within regions of the sinonasal cavity and pulmonary alveoli, sites of presumptive viral transmission and severe disease development for COVID-19, respectively. We show age, sex, and comorbidities do not increase ACE2 protein expression in the human respiratory tract.

*Implications of all the available evidence:* ACE2 protein abundance does not correlate with risk factors for severe clinical outcomes, but in some cases showed an inversed relationship. Features driving COVID-19 susceptibility and severity are complex, our data suggests factors other than ACE2 protein abundance as important determinants of clinical outcomes.

## Introduction

Angiotensin-converting enzyme 2 (ACE2) is the cellular receptor for both severe acute respiratory syndrome coronavirus (SARS-CoV) and SARS-CoV-2 (1, 2). SARS-CoV caused a pneumonia outbreak in 2002-2003 with a mortality rate of 9·6% and over 800 deaths worldwide (3). SARS-CoV-2 is the etiologic agent of coronavirus disease 2019 (COVID-19) which was first recognized in December 2019 and has now reached pandemic proportions (2, 4). SARS-CoV-2 infection can be fatal, with the risk for increased disease severity correlating with advanced age and underlying comorbidities, while children and younger individuals generally have milder disease (5–11). These trends in disease severity could reflect differences in ACE2 distribution and expression in the respiratory tract.

Previous studies have evaluated ACE2 expression in the respiratory tract. Studies of ACE2 mRNA transcript abundance have provided conflicting interpretations, as well as a lack of protein validation (12–19). ACE2 protein studies have been limited by a paucity of lung tissue substrates and reports that have yielded contradictory results (20–23) (Supplemental Table 1). It is reported that some clinical factors (sex, age, or presence of comorbidities) could influence ACE2 expression in the human lower respiratory tract. The ACE2 gene resides on the X chromosome and therefore could be differentially regulated between males and females due to variable X-inactivation (24). Increased abundance of circulating ACE2 protein is reported to correlate with male sex, advanced age, and chronic comorbidities such as diabetes, cardiovascular disease, and renal disease (reviewed in (25)). Recent single-cell mRNA sequencing (scRNA-seq) studies of respiratory tract cells have reported contradictory evidence regarding the correlation between ACE2 transcript abundance and age, sex, smoking status, and other comorbidities (12–17).

We investigated the hypothesis that ACE2 drives disease severity in susceptible patient populations through enhanced abundance or distribution in different locations or cell types of the respiratory tract. We reanalyzed publicly available scRNA-seq data from distal lung biopsies (26), nasal brushings, and nasal turbinate samples (27) to evaluate ACE2 transcript abundance in specific cell types. We complemented these analyses with optimized and validated ACE2 immunostaining protocols, to corroborate single cell analyses as well as to screen for differences in cellular ACE2 protein in lung tissues derived from a cohort of control and chronic diseased patients.

## Materials and methods

### Ethics

Studies on human tissues were approved by the institutional review board of the University of Iowa (Iowa IRB #199507432). Informed consent was obtained for all the tissues included in the study.

### Tissues

Tissues included nasal biopsies (n=3, deidentified and lacked evidence of significant disease or cancer), lung donors (n=29), primary cell cultures (28), and autopsy tissues (control tissues such as small intestine and kidney) that were selected from archival repositories as formalin-fixed paraffin-embedded blocks. All lungs were derived either from living donors at lung transplant or from deceased individuals maintained on life support for organ donation. In either case, lung tissues were routinely harvested according to transplant guidelines to maintain tissue viability. Lungs were surgically resected, placed in chilled media and transported to lab for examination, sample collection and fixation. Lung cases were selected to comprise two case study groups: 1) Chronic disease group was defined as having chronic comorbidities including: asthma, cardiovascular disease, chronic obstructive pulmonary disease, cystic fibrosis, diabetes, and smoking. 2) Control group was defined as lacking these chronic comorbidities and lacking clinical lung disease. The definition of chronic comorbidities was informed by reported independent risk factors for mortality in COVID-19 (8–10). The cumulative cohort included 29 cases (15 chronic comorbidities and 14 controls) with a broad range of ages (0·5 – 71 years) and both sexes were represented (13 female and 16 male). For these lungs, if a trachea or bronchus tissue block was available from the same case – these were included as well (Supplemental Table 2). Bronchioles were observed in most lung sections and were defined as intrapulmonary airways lacking evidence of cartilage or submucosal glands (29). Detailed medical histories other than diagnoses, including medication history, were not available.

### Immunohistochemistry and immunofluorescence

All formalin-fixed paraffin-embedded tissues were sectioned (~4 μm) and hydrated through a series of xylene and alcohol baths to water. Immunohistochemical techniques were used for the following markers: angiotensin-converting enzyme 2 (ACE2) (30), surfactant protein C (SP-C) (31), and mucin 5B (MUC5B) (32). For more specifics about the reagents please see Supplemental Table 3.

The immunostaining protocols for ACE2 were rigorously optimized and validated to avoid nonspecific staining that is commonplace and give confidence in the sensitivity of the protocol and quality of the tissues (Supplemental Figure 4, Supplemental Table 1 and Supplemental Table 3). We analyzed ACE2 protein expression in human upper and lower respiratory tract by immunohistochemistry (Supplemental Table 2). Human respiratory tract tissues were scored for ACE2 expression by a masked pathologist, following principles for reproducible tissue scores (17).

For immunofluorescence, formalin-fixed and paraffin-embedded human lung blocks were sectioned (~4 μm). Slides were baked (55°C × 15 min) and then deparaffinized (hydrated) in a series of xylene and progressive alcohol baths. Antigen retrieval was performed using Antigen Unmasking Solution (1:100, #H-3300) in citrate buffer (pH 6·0) solution to induce epitope retrieval (5 min × 3 times) in the microwave. Slides were washed (PBS, 3 times, 5 min each) and a PAP pen used to encircle the tissue. Slides were blocked with background blocking solution (2% BSA in Superblock 1 hr in humid chamber). Primary antibodies anti-ACE2 (1:100, Mouse monoclonal, MAB933, R&D Systems, Minneapolis, MN USA) and anti-TMPRSS2 (1:200, Rabbit monoclonal, #ab92323, Abcam, Cambridge, MA USA) were diluted in blocking solution (2% BSA in Superblock overnight 4°C). Secondary antibodies anti-mouse Alexa568 (for ACE2) and anti-rabbit Alexa488 (for TMPRSS2) were applied at a concentration of 1:600 for 1 hour at room temperature. Slides were washed and mounted with Vectashield containing DAPI.

### Tissue scoring

Stained tissue sections were examined for ACE2 localization using a post-examination method for masking and scored by a masked pathologist following principles for reproducible tissue scores (33). The initial examination showed a low heterogenous incidence of ACE2 staining for various tissues, so the following ordinal scoring system was employed to quantify number of staining-positive cells: 0 = below the limit of detection; 1 = <1%; 2 = 1-33%; 3 = 34-66%; and 4 = >66% of cells. For these anatomic regions (e.g. airway or alveoli), cell counts for each tissue were made to know the population density per microscopic field to make reproducible interpretations. For determination of AT2 cell size, ACE2 and SP-C protein immunostaining were evaluated on the same lung tissue section for each case. A region of minimally diseased lung was examined and SP-C^+^ AT2 cells were measured for diameter in the plane perpendicular to the basement membrane. Similar measurements were then made for ACE2^+^/SP-C^+^ cells.

### Analysis of single cell RNA sequencing data

Single cell RNA sequencing data sets were accessed from Gene Expression Omnibus (GEO) series GSE121600 (27) and GSE122960 (26). We included samples in our analysis if over 75% cell barcodes had >3000 unique molecular identifiers (UMI) in order to allow accurate cell type calls and clustering. Within samples passing our inclusion criteria, we retained cell barcodes with >1000 UMI. For GSE121600, raw H5 files for bronchial biopsy (GSM3439925), nasal brushing (GSM3439926), and turbinate (GSM3439927) samples were downloaded. Within series GSE121600, sample GSM3439925 (bronchial biopsy), 82% cell barcodes had less than 3000 UMI, so the sample was excluded from analysis. The turbinate sample was a biopsy from a 30-year-old female and the nasal brushing was performed in the inferior turbinate of a 56-year-old healthy male donor. For GSE122960, filtered H5 files for eight lung transplant donor samples from lung parenchyma (GSM3489182, GSM3489185, GSM3489187, GSM3489189, GSM3489191, GSM3489193, GSM3489195, GSM3489197) were downloaded. The eight donors varied from 21-63 years of age (median age = 48) and were composed of five African American, one Asian, and two white donors, and 2 active, 1 former, and 5 never smokers. Gene count matrices from the eight donors were aggregated for analysis.

Gene-by-barcode count matrices were normalized, log-transformed, and scaled followed by dimension reduction using principal components analysis (PCA). Principal components were used to obtain uniform manifold approximation and projection (UMAP) visualizations, and cells were clustered using a shared nearest neighbor (SNN) approach with resolution parameter 0·4, giving 14 clusters for nasal brushing, 15 clusters for turbinate, and 28 clusters for lung parenchyma. Cell types associated with each cluster were identified by determining marker genes for each cluster and comparing the list of marker genes to known cell type markers (Supplemental Figure 5). All analyses were performed using R package Seurat version 3.1.1 (34). In the nasal brushing sample, we were unable to associate a cell type with one cluster containing 776 cells (16·5%) due to low UMIs, so these cells were discarded.

For the lung parenchyma data, gene expression in alveolar type II cells for a single donor was quantified by summing up gene counts for all alveolar type II cells and dividing by total UMIs for all alveolar type II cells to get normalized counts, followed by rescaling the normalized counts to obtain counts per million (CPM).

### Statistical analyses

Statistical analyses for group comparisons and tissue scoring data were performed using GraphPad Prism version 8 (GraphPad Software, La Jolla, CA USA). Mann Whitney U tests or T-tests were used for group comparisons as appropriate. ACE2 protein detection in different tissues was analyzed using the ordinal scoring system (0-4) and Cochran-Armitage test for trend.

## Results

In the alveoli, ACE2 transcripts were detected mostly in alveolar type II (AT2) cells (89·5% of all ACE2^+^ cells) (Figure 1a), but specifically within a subset of these cells (1·2% of AT2 cells) (Figure 1b, Supplemental Figure 1a-b). These data indicate ACE2 transcripts are uncommon in most alveolar cell types. Alveoli had apical ACE2 protein only in a small number (usually ~1% or less) of AT2 cells (Figure 1c), consistent with the scRNA-seq results. The identity of these cells was confirmed by co-staining for surfactant protein-C. These ACE2^+^ AT2 cells were observed within areas of alveolar collapse (Figures 1d-f) and had morphologic features of hyperplastic AT2 cells, being more plump and larger than ACE2^−^ AT2 cells in the same tissue section (Figure 1g). Interestingly, alveolar macrophages were negative for ACE2 protein staining by immunohistochemistry, despite previous reports of ACE2 protein in these cells (Supplemental Table 1). The lack of ACE2 transcripts in macrophages was also confirmed by scRNA-seq data that revealed ACE2 mRNA in only 0·1% of macrophages, monocytes, or dendritic cells (Supplemental Figure 1a-b). The concordance between scRNA-seq and immunohistochemistry results provides compelling evidence that ACE2 is primarily present in a subset of AT2 cells and that alveolar macrophages lack ACE2.

**Figure 1.**
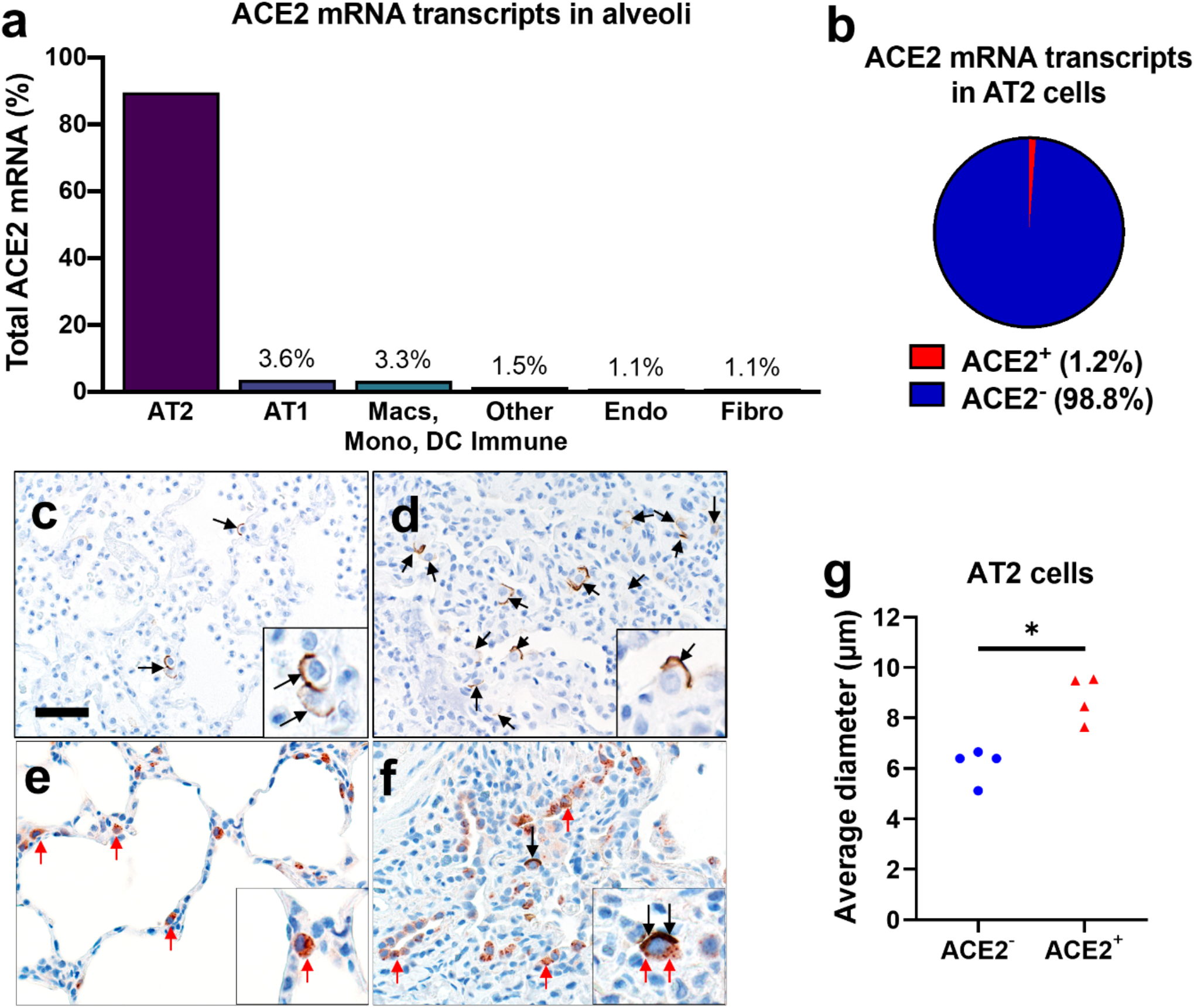
ACE2 expression in human lung. **a, b**) Single-cell RNA sequencing reanalysis of ACE2 transcript abundance in alveoli from lung parenchyma samples (26). Summative observations from all donors. Airway cells (basal, mitotic, ciliated, club) are not shown. **a**) 89·5% of the cells with detectable ACE2 mRNA in the alveoli are alveolar type II cells. **b**) Only 1·2% of alveolar type II cells have ACE2 mRNA transcripts. **c-f**) Detection of ACE2 protein (brown color, black arrows, and insets) in representative sections of lower respiratory tract regions and tissue scoring (see Supplemental Table 2) (**g**). **c, d**) Alveolar regions had uncommon to regional polarized apical staining of solitary epithelial cells (**c**) that (when present) were more readily detected in collapsed regions of lung (**d**). **e, f**) SP-C (red arrows, inset) and ACE2 (black arrows, inset) dual immunohistochemistry on the same tissue sections. **e**) Non-collapsed regions had normal SP-C^+^ AT2 cells lacking ACE2. **f**) Focal section of peri-airway remodeling and collapse with several SP-C^+^ (red arrows) AT2 cells, but only a small subset of AT2 cells had prominent apical ACE2 protein (black arrows, inset). **g**) SP-C^+^/ACE2^+^ AT2 cells were often larger than SP-C^+^/ACE2^−^ AT2 cells from same lung sections (see also **d** and **e** insets) indicative of AT2 hypertrophy, each data point represents the average value for each case from 5-10 cell measurements per group, P=0·0014, paired T-test. AT2: alveolar type II. AT1: alveolar type I. Macs: Macrophages. Mono: Monocytes. DC: dendritic cells. Other immune cells: B cells, mast cells, natural killer/T cells. Endo: Endothelial. Fibro: Fibroblasts/myofibroblasts. Bar = 35 μm.

Recent evidence indicates that proteases such as TMPRSS2 facilitate entry of SARS-CoV-2 into ACE2^+^ cells (35). We evaluated scRNA-seq data and observed that TMPRSS2 mRNA was present in 35·5% of all AT2 cells (Figure 2a) but was more prevalent (50·0%) in ACE2^+^ AT2 cells (Figure 2b). Additionally, we observed colocalization of ACE2 and TMPRSS2 on the apical membrane of these AT2 cells (Figure 2c). These findings suggest that AT2 cells with apical ACE2 and TMPRSS2 could readily facilitate SARS-CoV-2 cellular infection and disease as seen in COVID-19 patients.

**Figure 2.**
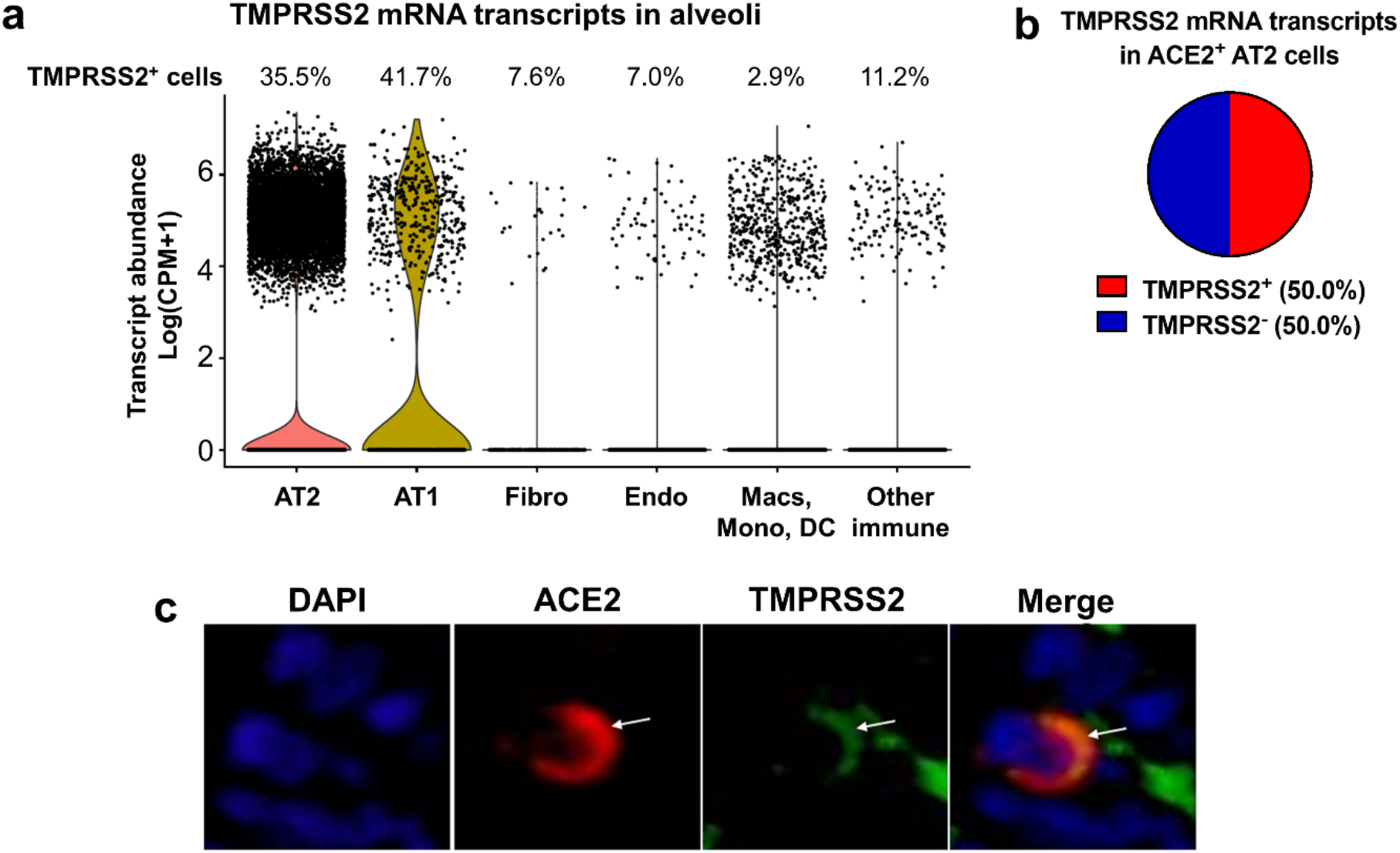
TMPRSS2 expression in the alveoli. **a, b**) Single-cell RNA sequencing reanalyses of TMPRSS2 transcript abundance in alveoli from lung parenchyma (26). Summative observations from all donors. **a**) Percentage of TMPRSS2^+^ cells within each cell type shows TMPRSS2 transcripts in 35·5% of alveolar type II cells. Airway cells (basal, mitotic, ciliated, club) are not shown. Violin plots represent expression, each data point denotes a cell. **b**) TMPRSS2 transcripts in ACE2^+^ alveolar type II cells. **c**) Immunofluorescence of alveoli shows apical colocalization of ACE2 and TMPRSS2 (white arrows). AT2: alveolar type II. AT1: alveolar type I. Macs: Macrophages. Mono: Monocytes. DC: dendritic cells. Other immune cells: B cells, mast cells, natural killer/T cells. Endo: Endothelial. Fibro: Fibroblasts/myofibroblasts. CPM: counts per million.

We next evaluated ACE2 in the conducting airways (trachea, bronchi, bronchioles). In the trachea and bronchi, apical ACE2 was rare and limited to ciliated cells (Figure 3a), similar to localization results in primary cultures of well-differentiated human airway epithelial cells (30). In the submucosal glands of large airways, occasional serous cells and vessels near the acini were positive for ACE2 (Supplemental Figures 2a-b). In bronchioles, ACE2 was regionally localized (Figures 3b-d). These findings show nominal detection of ACE2, corresponding with the lack of primary airway disease (e.g., bronchitis, etc.) seen in COVID-19 patients.

**Figure 3.**
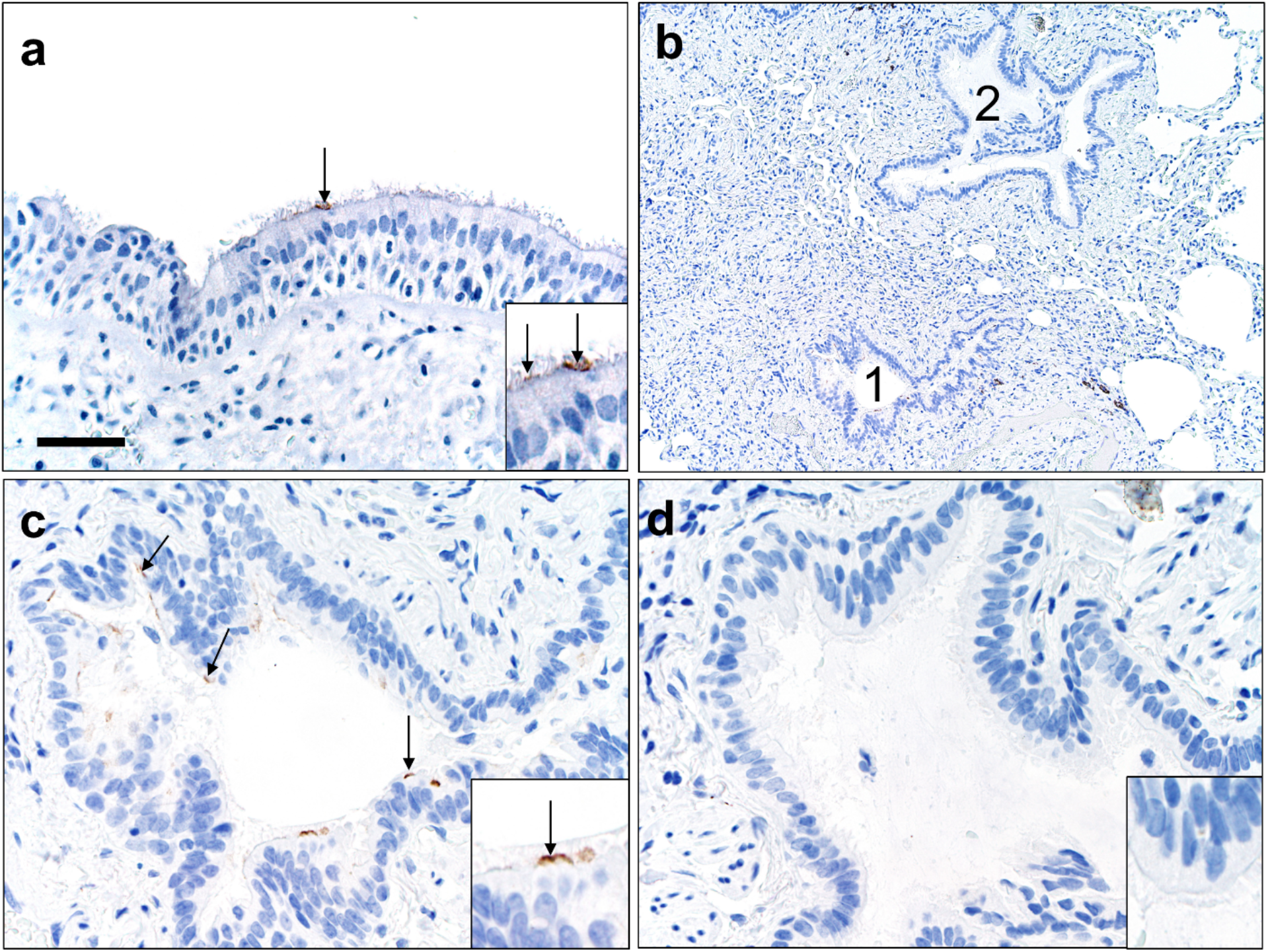
ACE2 protein in human lower airways. **a**) Large airways (trachea and bronchi) exhibited rare ACE2 protein on the apical surface of ciliated cells. **b-d**) Small airways (bronchioles) exhibited uncommon to localized apical ACE2 protein in ciliated cells (**c**, #1 in **b**) while the adjacent bronchioles (**d**, #2 in **b**) lacked protein. Bar = 35 (a), 140 (b), and 70 μm (c, d).

Detection of ACE2 protein has been variably reported in several small studies (21, 23). In this larger study, we saw that the regional distribution of ACE2 protein varied between donors. In the surface epithelium of trachea and bronchi, we detected ACE2 in only 12% and 27% of donors, respectively (Figure 4a). In the distal areas of the lung, ACE2 detection was more common, with bronchiolar and alveolar protein detection in 36% and 59% of donors, respectively (Figure 4a). A similar pattern of variable alveolar ACE2 was seen for mRNA transcripts in the scRNA-seq data, where 50% of donors showed lower abudance in AT2 cells, and the other 50% of donors showed higher abundance in the same cell type (Figure 4b). These findings suggest that ACE2 expression can vary between different lung regions and between individuals. Importantly, this low level of cellular protein provided us with an opportunity to investigate the potential for various clinical factors to increase ACE2 expression.

**Figure 4.**
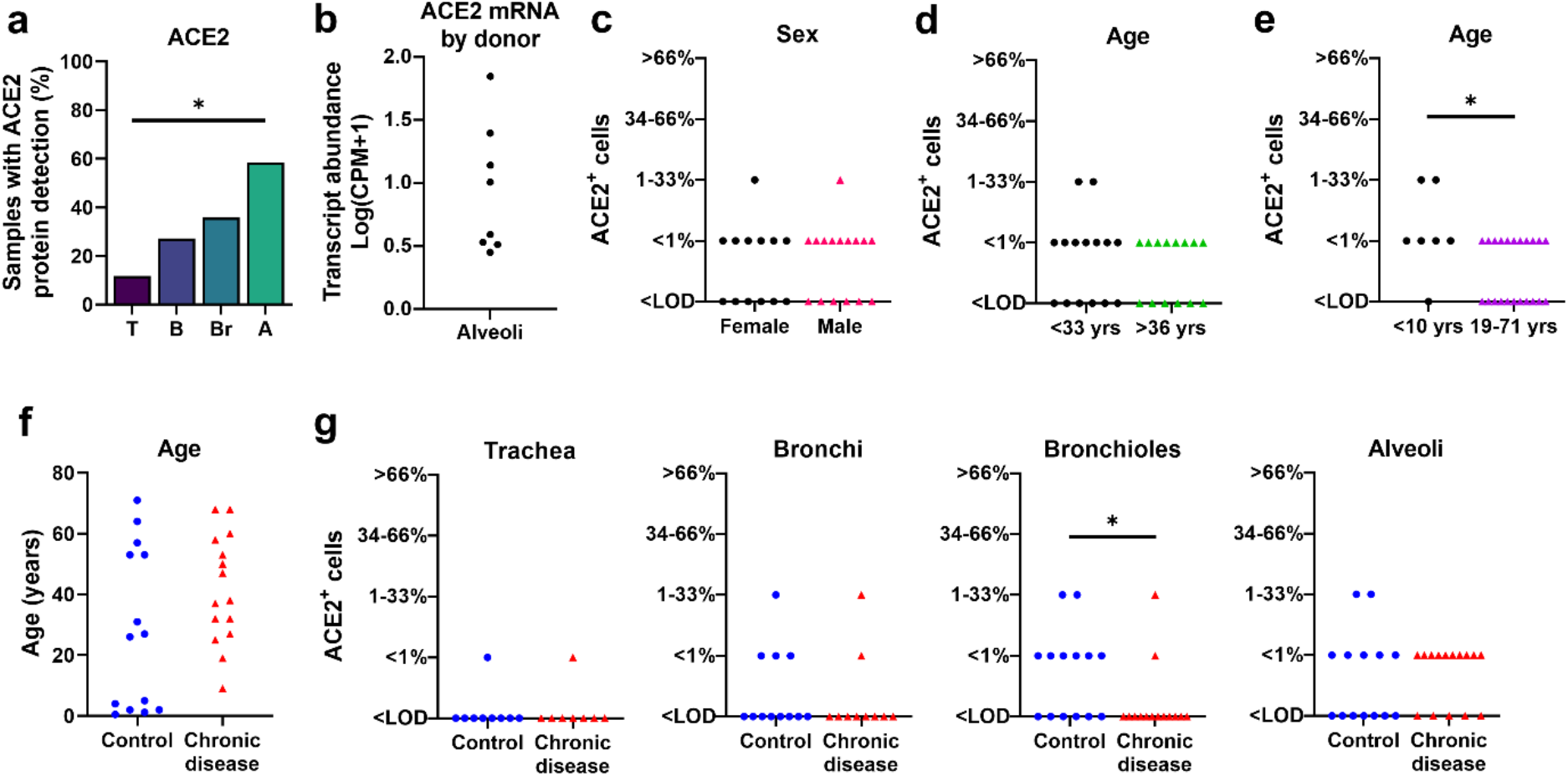
ACE2 distribution and scores in respiratory tissues. **a**) ACE2 protein had progressively increased detection between donors in tissues from trachea (T), bronchi (B), bronchioles (Br), to alveoli (A), (P=0·0009, Cochran-Armitage test for trend). **b**) Single-cell RNA sequencing reanalyses of ACE2 mRNA transcript abundance in the alveoli shows variation between donors (26). **c-d**) ACE2 protein scores from lung donors showed no differences based on sex or lower vs. upper ages (using median age as a cut-off) (A, P=0·7338 and B, P=0·7053, Mann-Whitney U test). **e**) ACE2 protein scores were elevated in young children (<10 yrs) compared to the remaining subjects (19-71 yrs) (P=0·0282 Mann-Whitney U test). **f**) Control and chronic disease groups did not have any significant differences in age (P=0·1362 Mann-Whitney U test). **g**) ACE2 protein scores for trachea, bronchi, bronchiole, and alveoli in control versus chronic disease groups (P= >0·9999, 0·6263, 0·0433, and 0·7359, respectively, Mann-Whitney U test). Each symbol represents one donor. CPM: counts per million. LOD: Limit of detection.

Independent risk factors associated with severe COVID-19 include male sex, increased age, and presence of comorbidities (6, 8-10, 36-38). To evaluate whether the spatial distribution and abundance of ACE2 protein in the lower respiratory tract differed by these risk factors, we scored tissues for ACE2 protein detection (Supplemental Table 2). In the cohort, neither age nor sex were associated with ACE2 protein detection (using the median age as cut-off) (Figures 4c-d). Since recent studies of COVID-19 infections suggested that young children have reduced disease severity when infected by SARS-CoV-2 (6, 7), we compared lung tissue samples from children <10 years of age to those from the remaining older subjects (19-71 years of age) and found that ACE2 protein detection was higher in this subset of young children (Figure 4e). To test whether ACE2 distribution was affected by the presence of underlying diseases, we assessed the ACE2 localization pattern using tissues from subjects with chronic comorbidities (asthma, cardiovascular disease, chronic obstructive pulmonary disease, cystic fibrosis, diabetes, and smoking) and compared them to controls (Supplemental Table 2). The control group was similar in age to the chronic disease group (Figure 4f). We observed no significant differences between the two groups in ACE2 distribution, except for bronchioles, where ACE2 protein was reduced in the chronic disease group (Figure 4g, Supplemental Figure 2c). These results show that ACE2 levels in the respiratory tract were not increased in association with risk factors for severe COVID-19, such as male sex, advanced age, and underlying chronic comorbidities. Instead, we saw increased ACE2 detection in children <10 years of age and in the small airways (bronchioles) of individuals without chronic comorbidities in our cohort, both groups with a lower risk for severe clinical disease.

Given the unexpected heterogeneity in the lower respiratory tract, we also investigated ACE2 in the upper respiratory tract. scRNA-seq data from nasal brushing and nasal turbinate samples (27) show ACE2 mRNA transcripts in 2-6% of epithelial cells (Supplemental Figure 3a-d). We then studied nasal biopsy tissues and found that ACE2 protein was detected in all tissue samples and, when present, was seen exclusively on the apical surface of ciliated cells. Distribution varied regionally based on the characteristics of the epithelium, with rare detection in thicker ciliated pseudostratified epithelium, and more abundant protein in thinner epithelium (Figures 5a-g). Thinner epithelial height is expected in specific regions including the floor of the nasal cavity, meatuses, and paranasal sinuses (39). The sinonasal cavity is an interface between the respiratory tract and the environment, and high SARS-CoV-2 viral loads can be detected in nasal swabs from infected patients (40), consistent with our ACE2 expression data. This reservoir of ACE2^+^ cells may facilitate the reported transmission from individuals who have very mild or asymptomatic disease (41).

**Figure 5.**
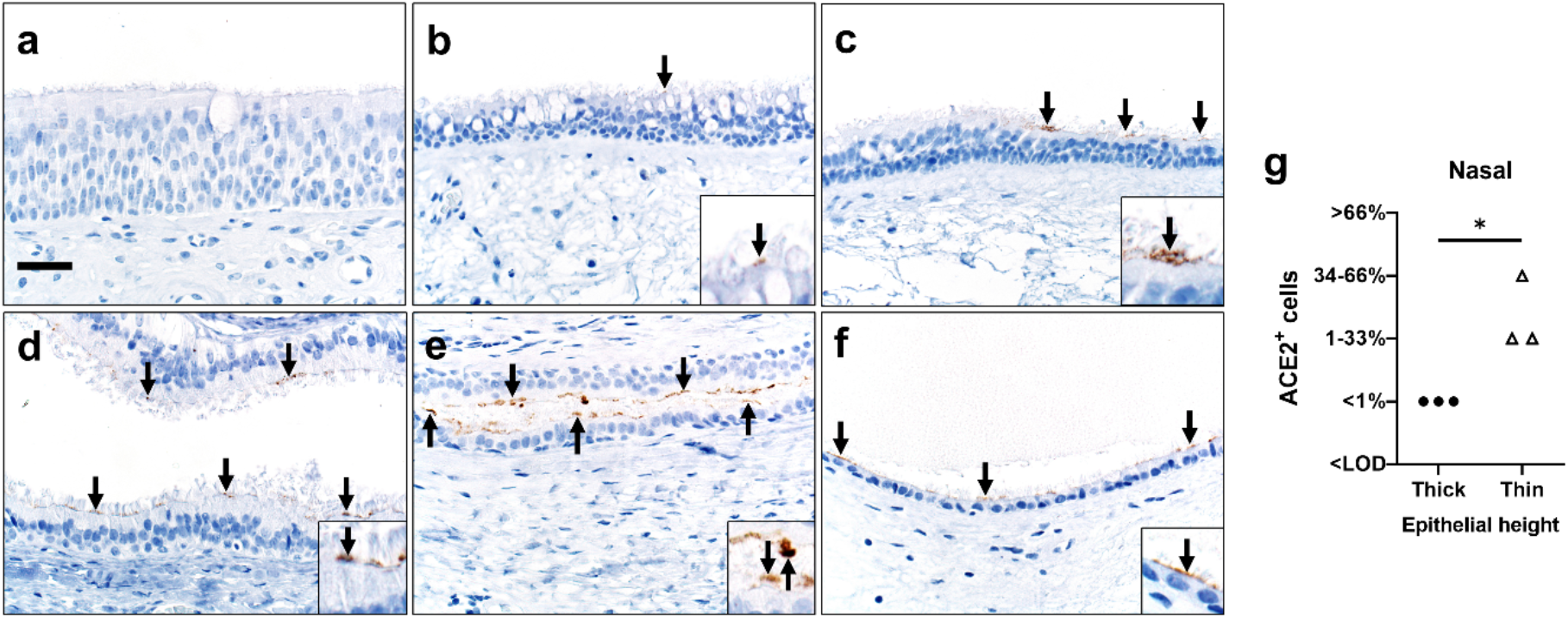
ACE2 protein in sinonasal tissues. Detection of ACE2 protein (brown color, arrows, and insets, **a-f**) and tissue scoring (**g**) in representative sections of nasal tissues. **a, b**) In thick pseudostratified epithelium (PSE) ACE2 protein was absent (**a**) to rare (**b**) and apically located on ciliated cells. **c**) Tissue section shows a transition zone from thick (left side, > ~4 nuclei) to thin (right side, ≤ ~4 nuclei) PSE and ACE2 protein was restricted to the apical surface of the thin PSE. **d-f**) ACE2 protein was detected multifocally on the apical surface of ciliated cells in varying types of thin PSE, even to simple cuboidal epithelium (**f**). Bar = 30 μm. **g**) ACE2 protein detection scores for each subject were higher in thin than thick epithelium, (P=0·05, Mann-Whitney U test). LOD: Limit of detection.

## Discussion

A critical aspect of this study was to evaluate ACE2 protein expression and distribution by immunohistochemistry to more accurately corroborate single cell transcript studies and better evaluate clinical groups for COVID-19 disease susceptibility. Previously, limited reports have variably shown ACE2 protein in the upper and lower respiratory tract, but cellular localization and distribution in human lung tissues have been inconsistent and contradictory (20–23) (Supplemental Table 1). *In vitro* studies demonstrate that ACE2 protein is found at the apical membrane of polarized airway epithelia, where it permits virus binding and cell entry (21, 30). In our study, ACE2 was consistently localized to the apical membranes of cells. ACE2 was more commonly found in the sinonasal cavity where transmission likely occurs and on AT2 cells of the lung parenchyma where severe disease develops. We sepeculate that expression of ACE2 in regions of the sinonasal cavity could explain the high transmissibility of SARS-CoV, SARS-CoV-2, and HCoV-NL63, a cold-related coronavirus which also uses ACE2 as a receptor.

SARS-CoV and SARS-CoV-2 both replicate in the lungs (42, 43), consistent with the ACE2 protein distribution defined in this study and suggested by previous studies (20, 21). We show that ACE2 and TMPRSS2 coexpress in AT2 cells at the mRNA and protein levels, suggesting susceptibility to infection. Additionally, it may also be possible that TMPRSS2^−^ ACE2^+^ AT2 cells can become infected through the use of other airway proteases (44). AT2 cells are critical for surfactant protein production and serve as progenitor cells for the AT1 cells, thus damage to these AT2 cells could contribute to acute lung injury (45), which is a common feature of severe COVID-19 (5). Additionally, the larger morphology of ACE2^+^ AT2 cells is consistent with a type of hyperplastic AT2 population that, if damaged, could affect the repair mechanisms of the alveoli. Infection of AT2 cells could disrupt epithelial integrity leading to alveolar edema, and facilitate viral spread to ACE2^+^ interstitial cells/vessels for systemic virus dissemination, given that SARS-CoV-2 has been detected in pulmonary endothelium (46) and blood (47). Furthermore, cell-to-cell spread of coronaviruses to other epithelial cells after initial infection could also occur via receptor-independent mechanisms related to the fusogenic properties of the S protein (48). It is interesting that computerized tomography studies of early disease in people with COVID-19 demonstrate patchy ground glass opacities in the peripheral and posterior lungs, regions that are more susceptible to alveolar collapse (49).

ACE2 protein detection in the lower respiratory tract was heterogeneous. The relatively small number of ACE2^+^ cells found in our study proved advantageous in evaluating whether conditions that predispose to severe disease also increased cellular ACE2 expression, but this was not observed. Rather we saw elevated ACE2 protein in demographic pools with expected low risk for severe COVID-19 (young children and in bronchioles of the control group) and these results suggest alternative explanations. First, the potential relationship between ACE2 abundance in the respiratory tract and severe COVID-19 is likely complex. On one hand, more receptor availability could enhance viral entry into cells and worsen disease outcomes; alternatively, ACE2 may play a protective role in acute lung injury through its enzymatic activity (50–52) and therefore could improve disease outcomes. Our data would support the latter and implicate a dualistic role for ACE2 as both a viral receptor and a protective agent in acute lung injury. Additionally, ACE2 exists in cell-associated and soluble forms (53). It is possible that greater ACE2 expression could result in increased soluble ACE2 in respiratory secretions where it might act as a decoy receptor and reduce virus entry (1, 54). Second, other factors such as TMPRSS2 expression might be more important in regulating disease severity. TMPRSS2 on the apical membrane of AT2 cells might facilitate SARS-CoV-2 entry when ACE2 is rare or even below the limit of detection in this study. Third, low levels of the receptor could be sufficient for the virus to infect and cause severe disease. Importantly, unlike SARS or HCoV-NL63, the SARS-CoV-2 spike glycoproteins undergo proteolytic processing at a multibasic S1/S2 site by furin intracellularly, prior to virion release (35, 55). Additionally, compared to SARS-CoV, the SARS-CoV-2 receptor binding motif has a higher affinity for ACE2 (56, 57). These features may enhance the ability of SARS-CoV-2 to bind to cells, undergo S2’ cleavage by TMPRSS2 or other surface proteases, fuse to the host cell membrane, and release its genome. Finally, we acknowledge that it is possible that SARS-CoV-2 infection could modify ACE2 expression in the respiratory tract, or that ACE2 expression in other organs could impact disease severity. It is important to mention that the lack of correlation between SARS-CoV-2 receptor expression and disease severity contrasts with another severe coronavirus disease, MERS, where comorbidities were observed to increase its receptor detection in respiratory tissues (58, 59).

mRNA transcript abundance is not always representative of protein levels (60), and therefore both should be evaluated in conjunction before making conclusions about gene expression. Some of the factors that account for these differences include post-transcriptional regulation or rapid protein turnover. Additionally, other factors limit direct comparisons between scRNA-seq results and protein staining, including sample size, tissue heterogeneity, and undefined biopsy sites. In the alveoli, we show ACE2 protein in a small subset of AT2 cells, which correlates with the scRNA-seq data and with other RNA sequencing publications (14, 18, 19). In the lower airways and sinonasal cavity, RNA sequencing data indicate ACE2 transcripts in both ciliated and secretory cells (14, 18, 19), but we show ACE2 protein is only found in ciliated cells. Likewise, some authors have reported lower ACE2 transcript abundance in children (12, 15) and suggested this finding as an explanation for the lower disease severity in this age group. In contrast, we show that children do not have less ACE2 protein than older adults, and while children appear protected from severe lung disease, they are likely at similar risk for infection (7, 61, 62).

In summary, we find that ACE2 protein has heterogeneous expression in the respiratory tract with more frequent ACE2 detection in the sinonasal epithelium and AT2 cells that correlates with putative sites for transmission and severe disease, respectively. The small subset of ACE2^+^ AT2 cells in the lung could be further studied to reveal factors regulating ACE2 expression and clarify potential targets for antiviral therapies. Contrary to our initial hypothesis, we saw no increase of ACE2 protein in the chronic disease group. Interestingly, we observed increased ACE2 in young children and control group bronchioles, suggesting a possible protective effect by ACE2 expression. These results suggest that features driving disease susceptibility and severity are complex. Factors other than ACE2 protein abundance, including viral load, host innate and adaptive immune responses, and the activities of the pulmonary renin-angiotensin system may also be important determinants of outcomes.

## Author contributions

Conceptualization and writing – original draft, M. E.O.B., P.B.M. and D.K.M.; Data curation, A.T.; Formal analysis, M. E.O.B., A.T., A. A.P. and D.K.M.; Investigation, A.T., A. A.P., M.R.L., C.W.-L. and D.K.M.; Visualization, M. E.O.B., A.T., D.K.M.; Resources, A. A.P., J.A.K.-T., P.H.K., P.T., P.B.M. and D.K.M.; Writing – review and editing, M. E.O.B, A.T., A.A.P., M.R.L., J.A.K.-T., P.H.K., P.T., C.W.-L., P.B.M. and D.K.M.

All authors approved the final version of this manuscript.

## Acknowledgements

We thank our laboratory members and colleagues Stanley Perlman, Robert Robinson, and Tom Gallagher for their helpful discussion and technical assistance.

## Data sharing statement

All single-cell RNA sequencing datasets used in this manuscript are publicly available as outlined in the Methods.

## Supplemental information

**Supplemental Figure 1.**
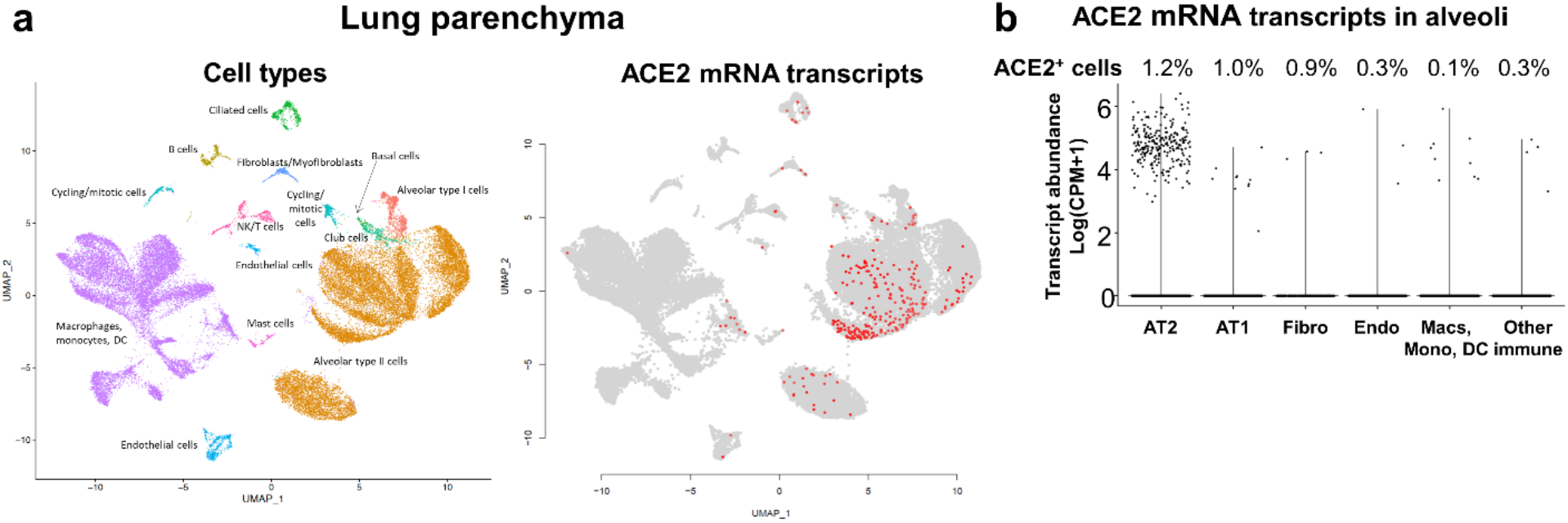
Single-cell RNA sequencing reanalyses of ACE2 transcript abundance in lung parenchyma (26). Summative observations from all donors. **a**) Uniform manifold approximation and projection (UMAP) visualizations. Cells were clustered using a shared nearest neighbor (SNN) approach. Cell types associated with each cluster were identified by determining marker genes for each cluster. Each data point denotes a cell. On the right panel, cells with ACE2 transcripts are shown in red. **b**) Violin plots representing ACE2 expression in the alveoli. Airway cells (basal, mitotic, ciliated, club) are not shown. Percentage of ACE2^+^ cells within each cell type shows ACE2 transcripts in 1·2% of alveolar type II cells and in 0·1% of macrophages, monocytes, or dendritic cells. Each data point denotes a cell, most cells have no expression (0). AT2: alveolar type II. AT1: alveolar type I. Macs: Macrophages. Mono: Monocytes. DC: dendritic cells. Other immune cells: B cells, mast cells, natural killer/T cells. Endo: Endothelial. Fibro: Fibroblasts/myofibroblasts. NK: Natural killer. CPM: Counts per million.

**Supplemental Figure 2.**
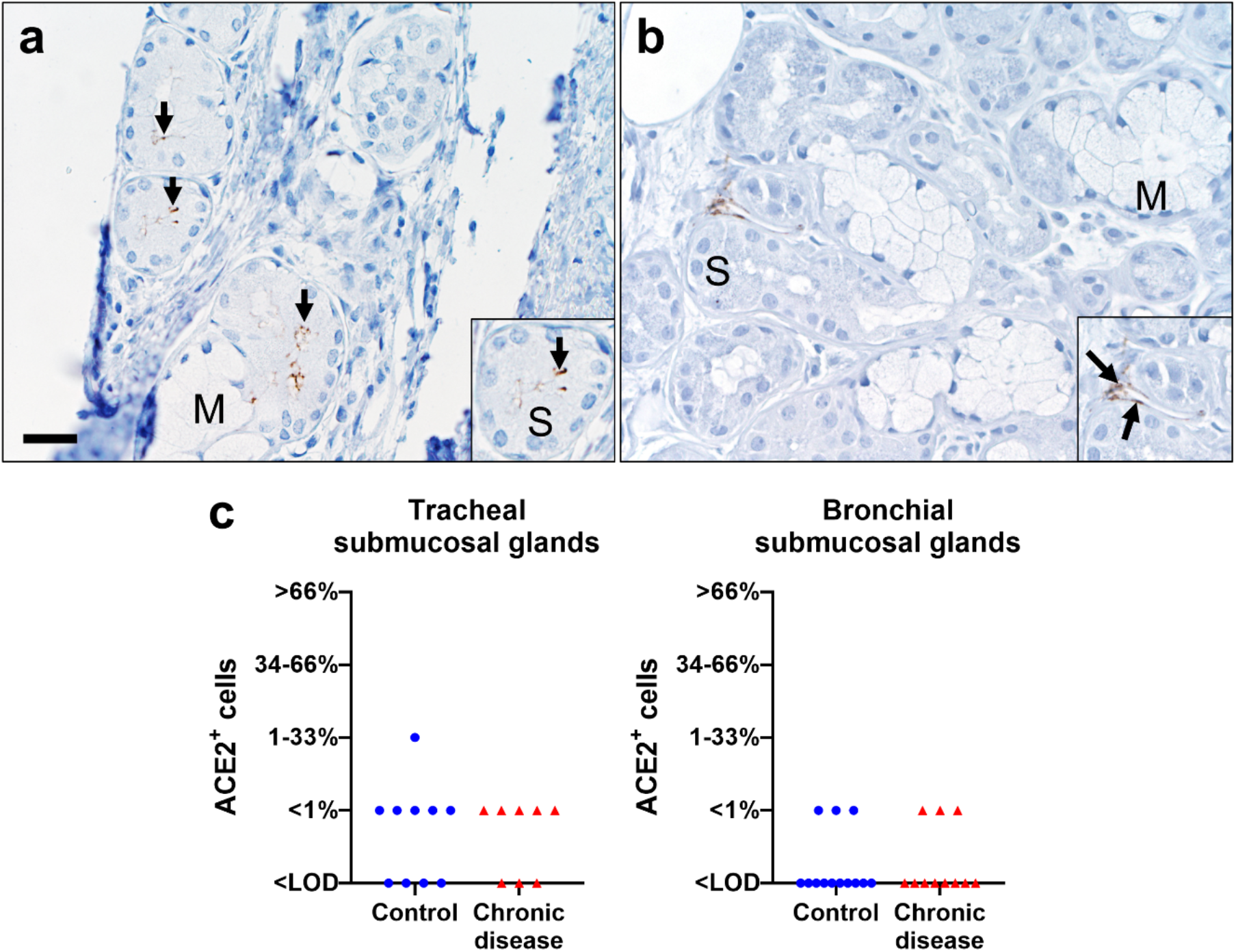
Representative tissue section from submucosa of large airways (trachea/bronchi) showing ACE2 protein localization (brown color, black arrows) (**a, b**) and scores (**c**). **a**) Submucosal glands had uncommon to localized apical ACE2 protein (arrows) in serous (S) cells, but not mucous (M) cells. **b**) Submucosal glands also had absent to uncommon ACE2 protein (arrows) in the interstitium that centered on vascular walls and endothelium. This vascular staining was uncommonly seen in lung too and corresponded to the low levels seen in transcripts for these endothelial cells (Supplemental Figure 1a-b). Note the absence of ACE2 staining in serous (S) or mucous (M) cells of the gland (**b**). **c**) ACE2 protein scores for each subject for serous cells in submucosal glands from trachea and bronchi, in control versus chronic disease groups (P>0·9999, 0·9999, respectively, Mann-Whitney U test). Bar = 25 μm. LOD: Limit of detection.

**Supplemental Figure 3.**
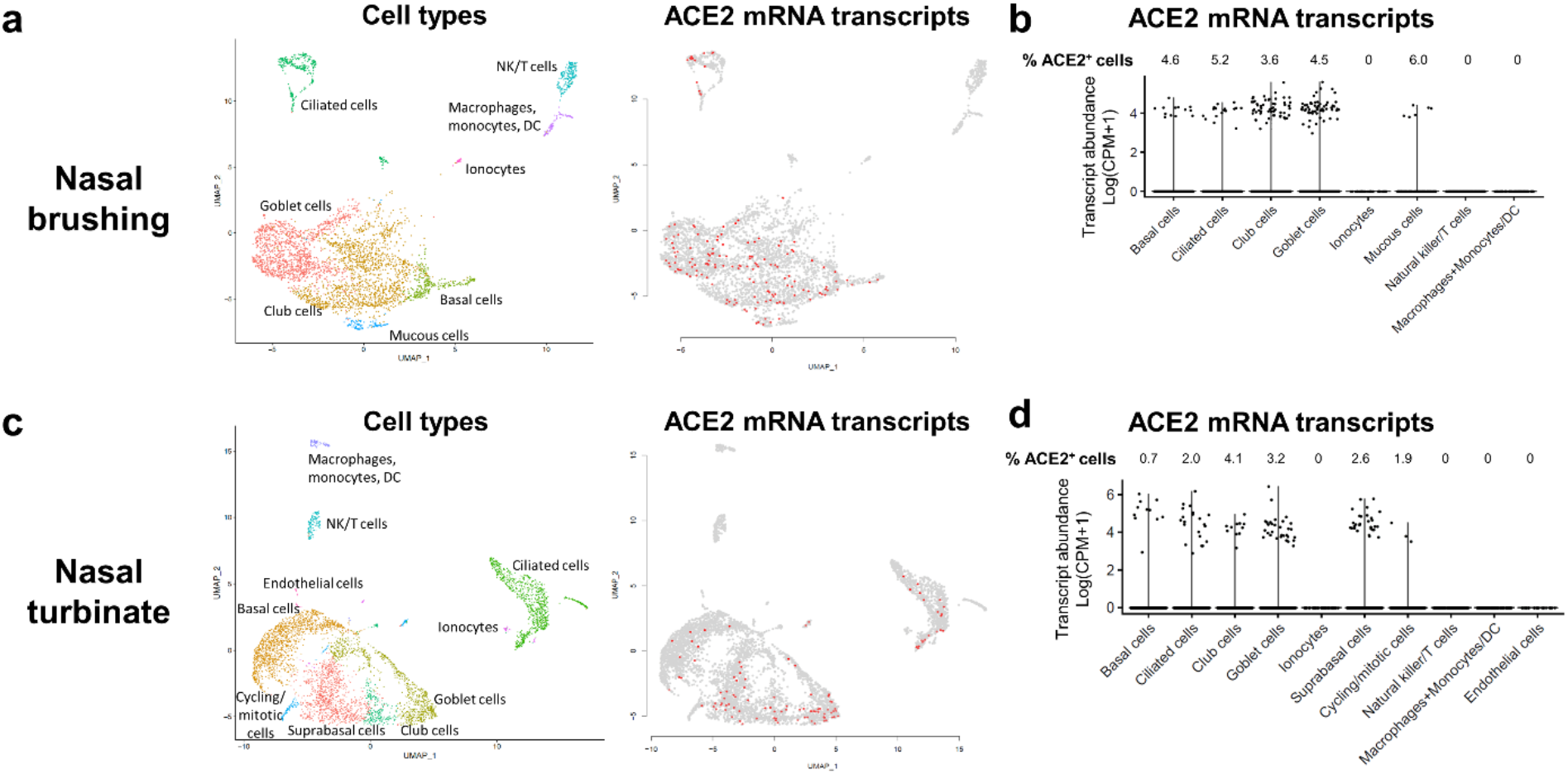
Single-cell RNA sequencing reanalyses of ACE2 transcript abundance in nasal brushing (**a, b**) and nasal turbinate (**c, d**) (27). **a, c**) Uniform manifold approximation and projection (UMAP) visualizations. Cells were clustered using a shared nearest neighbor (SNN) approach. Cell types associated with each cluster were identified by determining marker genes for each cluster. Each data point denotes a cell. On the right panels, cells with ACE2 transcripts are shown in red. **b, d**) Violin plots representing ACE2 expression. In nasal turbinate and nasal brushing, percentage of ACE2^+^ cells within each cell type shows ACE2 expression on epithelial cells. Each data point denotes a cell, most cells have no expression (0). DC: dendritic cells. NK: Natural killer. CPM: Counts per million.

**Supplemental Figure 4.**
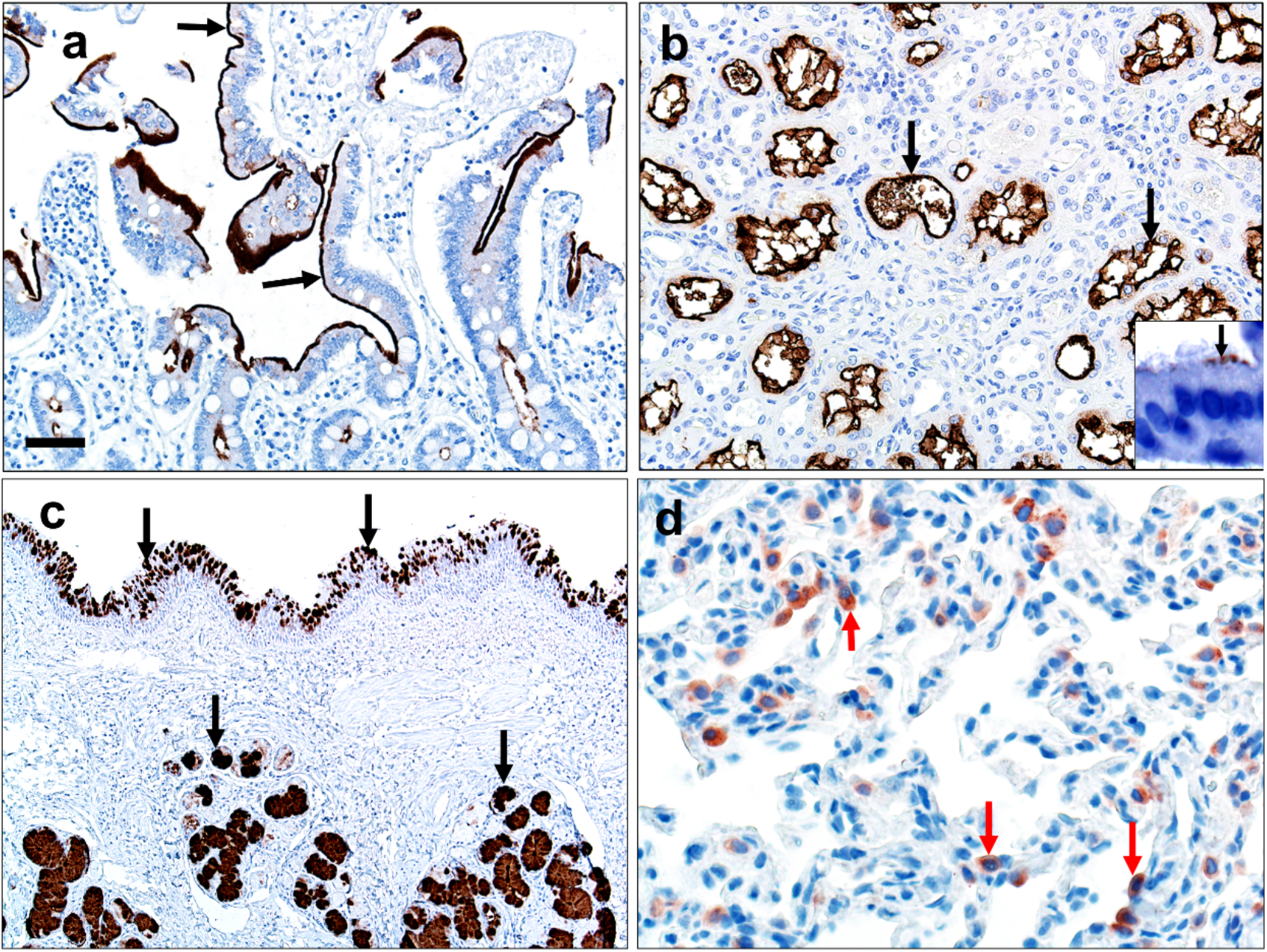
Quality controls for ACE2 immunohistochemistry technique (**a, b**) and tissue quality (**c, d**). **a, b**) ACE2 protein (brown color, black arrows) was detected along the apical surface of small intestine enterocytes (**a**), renal tubule epithelium (**b**), and ciliated cells (**b, inset**) of primary airway cell cultures. These findings demonstrate specific detection of ACE2 protein in cells/tissues consistent with known ACE2 expression. **c**) Representative immunostaining of bronchus detected abundant MUC5B protein (brown color, black arrows) in mucous cells of surface epithelium (top) and submucosal glands (bottom). **d**) Representative sections of alveoli had SP-C^+^ alveolar type II cells (red color, red arrows). These results (**c, d**) demonstrate the tissues were intact and that immunostaining can be used to detect native airway (**c**) and lung (**d**) proteins. Bar = 40 (a, b), 80 (c), and 20 μm (d).

**Supplemental Figure 5.**
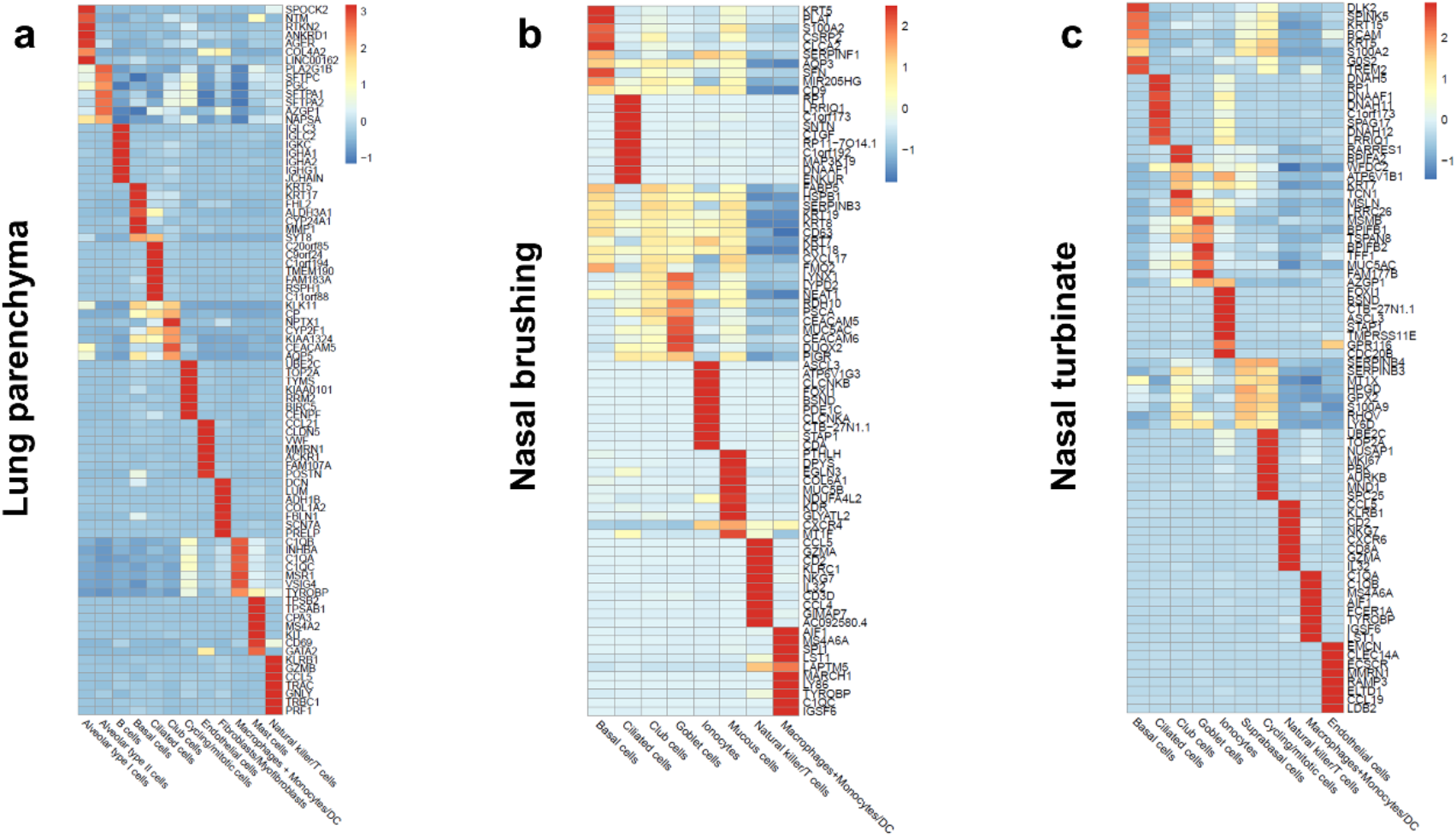
Single-cell RNA sequencing reanalyses of lung parenchyma (a) (26), nasal brushing (**b**), and nasal turbinate (**c**) (27). Heatmaps depicting the marker genes for each cluster that were used to assign cell types.

**Supplemental Table 1.**
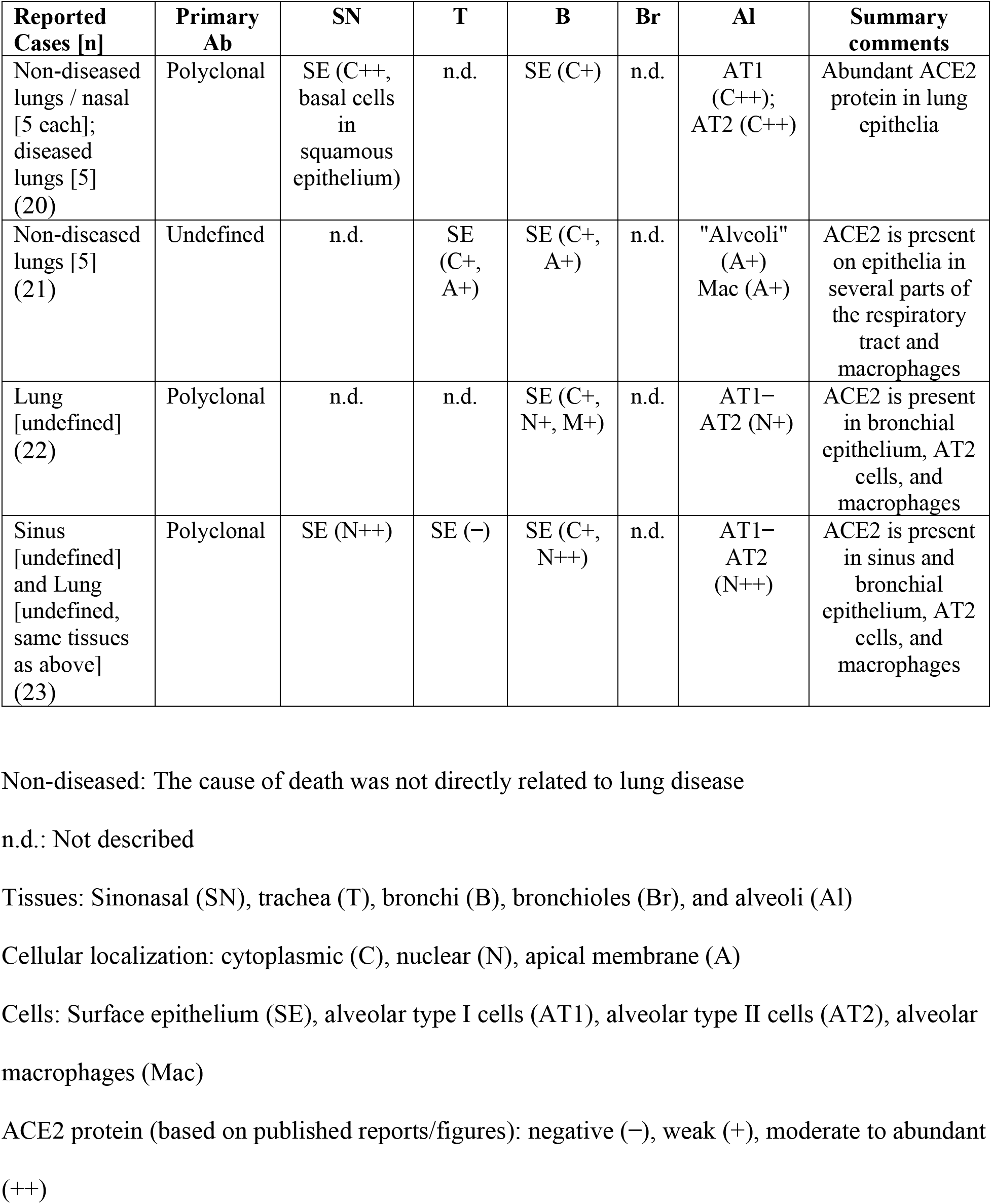
ACE2 protein reported in surface epithelium (SE) of human respiratory tract surface epithelium.

| **Reported Cases [n]** | **Primary Ab** | **SN** | **T** | **B** | **Br** | **Al** | **Summary comments** |
| --- | --- | --- | --- | --- | --- | --- | --- |
| Non-diseased lungs / nasal [5 each]; diseased lungs [5] (20) | Polyclonal | SE (C++, basal cells in squamous epithelium) | n.d. | SE (C+) | n.d. | AT1 (C++); AT2 (C++) | Abundant ACE2 protein in lung epithelia |
| Non-diseased lungs [5] (21) | Undefined | n.d. | SE (C+, A+) | SE (C+, A+) | n.d. | "Alveoli" (A+)<br>Mac (A+) | ACE2 is present on epithelia in several parts of the respiratory tract and macrophages |
| Lung [undefined] (22) | Polyclonal | n.d. | n.d. | SE (C+, N+, M+) | n.d. | AT1–AT2 (N+) | ACE2 is present in bronchial epithelium, AT2 cells, and macrophages |
| Sinus [undefined] and Lung [undefined, same tissues as above] (23) | Polyclonal | SE (N++) | SE (–) | SE (C+, N++) | n.d. | AT1–AT2 (N++) | ACE2 is present in sinus and bronchial epithelium, AT2 cells, and macrophages |

Non-diseased: The cause of death was not directly related to lung disease
n.d.: Not described
Tissues: Sinonasal (SN), trachea (T), bronchi (B), bronchioles (Br), and alveoli (Al)
Cellular localization: cytoplasmic (C), nuclear (N), apical membrane (A)
Cells: Surface epithelium (SE), alveolar type I cells (AT1), alveolar type II cells (AT2), alveolar macrophages (Mac)
ACE2 protein (based on published reports/figures): negative (–), weak (+), moderate to abundant (++)

**Supplemental Table 2.**
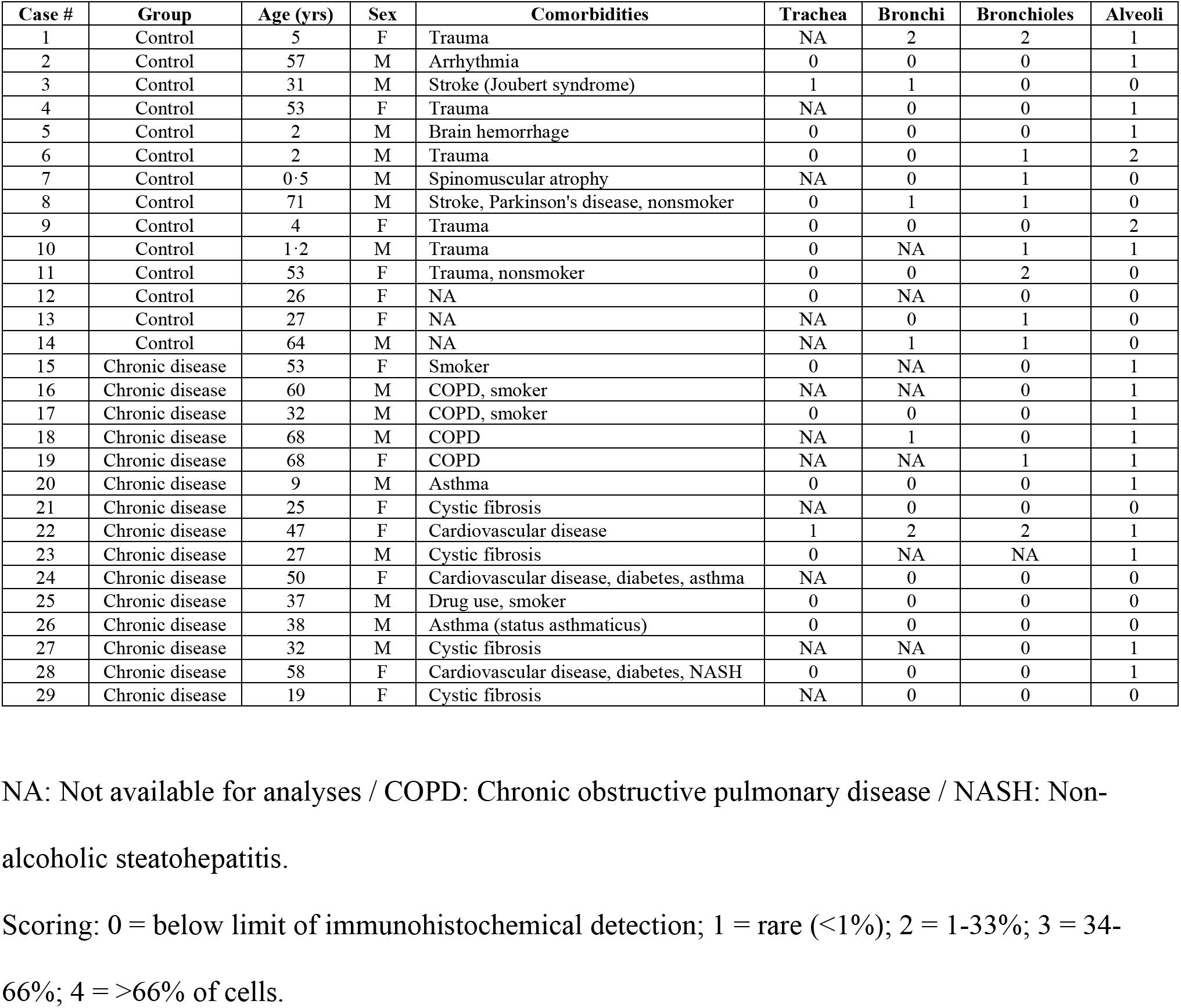
Donor demographics and ACE2 distribution scores for each tissue region.

**Supplemental Table 3.** Parameters for immunohistochemistry on fixed tissues.

| Target | Primary Antibody | Antigen Retrieval | Secondary Reagents |
| --- | --- | --- | --- |
| Angiotensin-<br>Converting<br>Enzyme 2 (ACE2) | Anti-ACE2,<br>monoclonal (MAB933,<br>R&D Systems,<br>Minneapolis, MN<br>USA) in diluent at<br>1:100 x 1 hour. | HIER, Citrate Buffer,<br>pH 6·0, 110°C for 15<br>minutes; 20 min cool<br>down (Decloaking<br>Chamber Plus, Biocare<br>Medical, Concord, CA<br>USA) | Dako EnVision+ System-<br>HRP Labeled Polymer<br>Anti-mouse, 60 min (Dako<br>North America, Inc.,<br>Carpentaria, CA USA),<br>DAB Chromogen,<br>counterstain. |
| MUC5B | Rabbit anti-MUC5B<br>polyclonal, (LSBio<br>#LS-B8121, LifeSpan<br>BioSciences, Inc.,<br>Seattle, WA) in Dako<br>Antibody Diluent<br>(Dako North America,<br>Inc., Carpentaria, CA);<br>1:60,000/30 min | HIER, Citrate buffer<br>pH 6·0, 110°C for<br>15min; 20 min cool<br>down | Step 1: Biotinylated anti-Rabbit IgG (H+L) (Vector<br>Laboratories, Inc.,<br>Burlingame, CA) in Dako<br>Wash Buffer (Dako North<br>America, Inc., Carpentaria,<br>CA); 1:500, 30 min<br>Step 2: Vectastain ABC<br>Kit (Vector Laboratories,<br>Inc., Burlingame, CA),<br>30min. DAB Chromogen,<br>counterstain. |
| Surfactant Protein<br>– C (SP-C) | Anti-SP-C, polyclonal<br>(PA5-71680, Thermo<br>Fisher Scientific,<br>Waltham, MA USA) in<br>diluent 1:100 x 1 hour | HIER, Citrate Buffer,<br>pH 6·0, 110°C for 15<br>minutes; 20 min cool<br>down (Decloaking<br>Chamber Plus, Biocare<br>Medical, Concord, CA<br>USA) | Dako EnVision+ System-<br>HRP Labeled Polymer<br>Anti-rabbit, 60 min (Dako<br>North America, Inc.,<br>Carpentaria, CA USA),<br>AEC chromogen,<br>counterstain. |

HIER – Heat-induced epitope retrieval
DAB – 3,3'-Diaminobenzidine (produces brown stain)
AEC - aminoethyl carbazole (produces red stain)
Counterstain – Harris hematoxylin (blue color)

